# Neoblast-like Stem Cells of *Fasciola hepatica*

**DOI:** 10.1101/2023.12.15.571799

**Authors:** Paul McCusker, Nathan G Clarke, Erica Gardiner, Rebecca Armstrong, Erin M McCammick, Paul McVeigh, Emily Robb, Duncan Wells, Madelyn Nowak-Roddy, Abdullah Albaqami, Angela Mousley, Jonathan A Coulter, John Harrington, Nikki Marks, Aaron Maule

## Abstract

The common liver fluke (*Fasciola hepatica*) causes the disease fasciolosis, which results in considerable losses within the global agri-food industry. There is a shortfall in the drugs that are effective against both the adult and juvenile life stages within the mammalian host, such that new drug targets are needed. Over the last decade the stem cells of parasitic flatworms have emerged as reservoirs of putative novel targets due to their role in development and homeostasis, including at host-parasite interfaces. Here, we investigate and characterise the proliferating cells that underpin development in *F. hepatica*. We provide evidence that these cells are capable of self-renewal, differentiation, and are sensitive to ionising radiation - all attributes of neoblasts in other flatworms. Changes in cell proliferation were also noted during the early stages of *in vitro* juvenile growth/development (around four to seven days post excystment), which coincided with a marked reduction in the nuclear area of proliferating cells. Furthermore, we generated transcriptomes from worms following irradiation-based ablation of neoblasts, identifying 124 significantly downregulated transcripts, including known stem cell markers such as *fgfrA* and *plk1*. Sixty-eight of these had homologues associated with neoblast-like cells in *Schistosoma mansoni*. Finally, RNA interference mediated knockdown of histone *h2b* (a marker of proliferating cells), ablated neoblast-like cells and impaired worm development *in vitro*. In summary, this work demonstrates that the proliferating cells of *F. hepatica* are equivalent to neoblasts of other flatworm species and demonstrate that they may serve as attractive targets for novel anthelmintics.

**Author Summary:** Liver fluke are parasitic worms that infect both livestock and humans worldwide, threatening food security and human health. Treatments against this disease-causing parasite are limited, and growing resistance to drugs is undermining the effectiveness of control strategies. Since drugs represent the only viable control option, it is crucial that new drugs are discovered through the identification and validation of new drug targets. Stem cells play important roles in the normal growth and repair processes of many organisms, but when these cells become dysregulated through mutation, they can drive the development of cancers. Stem cells of liver fluke may be attractive novel drug targets as disruption would affect worm survival and/or development within their host. In this research we describe the characteristics of liver fluke stem cells, such as their sensitivity to radiation and their ability to develop into new cell types (key stem cell features). We used radiation in combination with RNA sequencing to identify genes associated with the liver fluke stem cells. Finally, we used reverse genetics to reduce the expression of a gene associated with stem cells, which led to the loss of stem cells and reduced worm growth/development. These data provide evidence to support the exploitation of stem cells as a source of novel drug targets for liver fluke control.

## Introduction

Liver fluke of the genus *Fasciola* have a worldwide distribution and are known for their impact on both human and animal health. Within their definitive mammalian hosts *Fasciola* cause both acute and chronic disease. Losses due to liver fluke within the agri-food sector are conservatively estimated at >$3 billion/year [1]. Furthermore, ∼2.4 million humans suffer from fascioliasis, the human form of the disease, leading to its classification as a Neglected Tropical Disease [2]. The acute form of the disease is caused by juvenile worms burrowing through the liver parenchyma. After 3-10 weeks, depending on host, juveniles move into the bile ducts where they mature into egg-laying adults, initiating the chronic form of the disease. While a few drugs (e.g. closantel and nitroxynil) treat the adult fluke, treatment options for the migrating juveniles are limited. The drug of choice, triclabendazole (TCBZ), is the only flukicide effective at treating both acute and chronic infections [3]. However, resistance to TCBZ was first documented 30 years ago [4], and has since been observed on multiple continents [5]. No novel flukicides have been developed in recent years such that new drugs are needed for the sustainable control of fasciolosis, particularly the acute disease, as it threatens ruminant production systems. Moreover, the impact of fasciolosis on global food security is further complicated by climate change [6].

One potential source of novel drug targets within *Fasciola* are the somatic stem cells that drive growth and development. Stem cells are undifferentiated cells with the capacity to self-renew and potential to differentiate into specific cell types. Flatworm stem cells, known as neoblasts, have been of interest in free-living flatworms for over a century [7] due to the remarkable abilities of many planarians to regenerate tissue [8]. Platyhelminth neoblasts are pluripotent (capable of differentiating into multiple embryonic layers), constitute 20-35% of the cell population [9], and are the only actively proliferating somatic cells [10]. Several neoblast populations, and associated marker transcripts, have been characterised [11–15]. Clonogenic neoblasts (cNeoblasts) are one of the most remarkable subpopulations with the ability to recover all cell types in irradiated worms devoid of neoblasts, thereby rescuing worms which would otherwise die [16]. Stem cells are also susceptible to ablation via irradiation [17] such that it has become an indispensable tool for flatworm neoblast characterisation [18].

While older studies mention the presence of germinative/embryonic cells in parasitic flatworms [19–22], and even regenerative abilities in schistosomes [23], it was only within the last decade that research into parasitic flatworm stem cells has begun in earnest. Somatic neoblast-like stem cells were identified in different *Schistosoma mansoni* life stages [24,25], expressing known markers of the neoblasts found in their free-living cousins (e.g. *fgfrA* and *nanos2*). Since then, these neoblast-like cells have been found to be involved in wound repair [26], tegumental cell turnover [27,28] germline development [29], digestive tract development [30], and esophageal gland development [31] in *S. mansoni*. Germinative/stem cells in cestodes drive metacestode development in *Echinococcus granulosus* [32], and proglottid regeneration in *Hymenolepis diminuta* and *Mesocestoides corti* [33,34].

We previously described how proliferating neoblast-like cells drive growth/development in *in vitro F. hepatica* [35], finding this growth to be concurrent with the upregulation of known neoblast markers (*argonaute* and *nanos*). Additionally, we observed that treatment with the antimetabolite hydroxyurea inhibited both cell proliferation and worm growth. Further omics studies have shown how proliferation rates of *F. hepatica* neoblast-like cells are controlled by temperature [36], and that immature worms migrating through the host liver upregulate signal transduction pathways associated with stem cells, such as the PI3K/AKT/mTOR, AMPK and Hippo pathways [37]. A recent publication provided indirect transcriptomic evidence that *F. hepatica* neoblast-like cells express transcripts classically associated with neoblasts, where faster growing *in vivo* worms (in contrast to slower growing *in vitro* worms) upregulate numerous cell cycle associated genes including cyclins, minichromosome maintenance complex members and kinases, such as polo-like kinase 1 [38]. Curiously, this study also revealed that faster growing worms have a marked downregulation of transcripts related to the nervous system, suggesting a potential role for the nervous system in the regulation of stem cells/growth [38].

In this study we set out to further characterise the neoblast-like cells of *F. hepatica*, examining their dynamics in growing worms as well as their ability to self-renew and differentiate. Additionally, we evaluated their susceptibility to ablation via irradiation and used transcriptomics on irradiated worms to identify genes closely associated with *F. hepatica*’s neoblast-like cells. Finally, we used reverse genetics to silence a stem cell associated gene, histone *h2b*, examining the effects of knockdown on worm growth and development *in vitro*.

## Results and Discussion

### *F. hepatica* neoblast-like cell proliferation underpins *in vitro* growth

Our previous studies demonstrated how *F. hepatica* growth appears to be linked to proliferation of their neoblast-like cells [35,38]. We further explored this link, and the kinetics of the neoblast-like cells, by comparing proliferation rates of growing and non-growing worms across three weeks through 24-hour incubations in EdU, a thymidine analogue that labels cells going through the S phase of the cell cycle. We confirmed that growth was connected to proliferation rates, since an increase in the number of proliferating cells was observed in worms growing in CS50 (+CS), a trend not seen in non-growing worms maintained in RPMI alone (-CS) (Fig 1A). Significant differences were observed in the number of EdU^+^ nuclei at all timepoints examined (Fig 1B), even as early as two days post excystment (#EdU^+^ nuclei: -CS, 13±1; +CS, 29±1.5). This demonstrates how quickly juveniles respond to growth stimuli and is indicative of the extraordinary growth rates observed in *in vivo F. hepatica* juveniles [39]. By nine days post excystment +CS juveniles possessed >10x the number of neoblast-like cells that -CS juveniles did (#EdU^+^ nuclei: -CS, 22±1; +CS, 303±14), after which the viability of -CS juveniles deteriorated to the point that further comparisons via EdU incorporation assays were not informative. The link between growth and proliferative cells was further illustrated by plotting worm area against number of EdU+ nuclei, demonstrating a clear positive correlation (Fig 1C, R^2^=0.54, n=210, p<0.0001). This correlation has been noted previously in *Dugesia* where larger worms had more neoblasts, though not a greater density [40,41]. This correlation between size and neoblast-like cell proliferation was stronger than the relationship between neoblast-like cell proliferation and juvenile age (Fig 1C), suggesting that smaller worms are potentially feeding less. Indeed, studies in planarians have shown that lack of food results in a reduction in the number of neoblast progeny [42].

**Fig 1.**
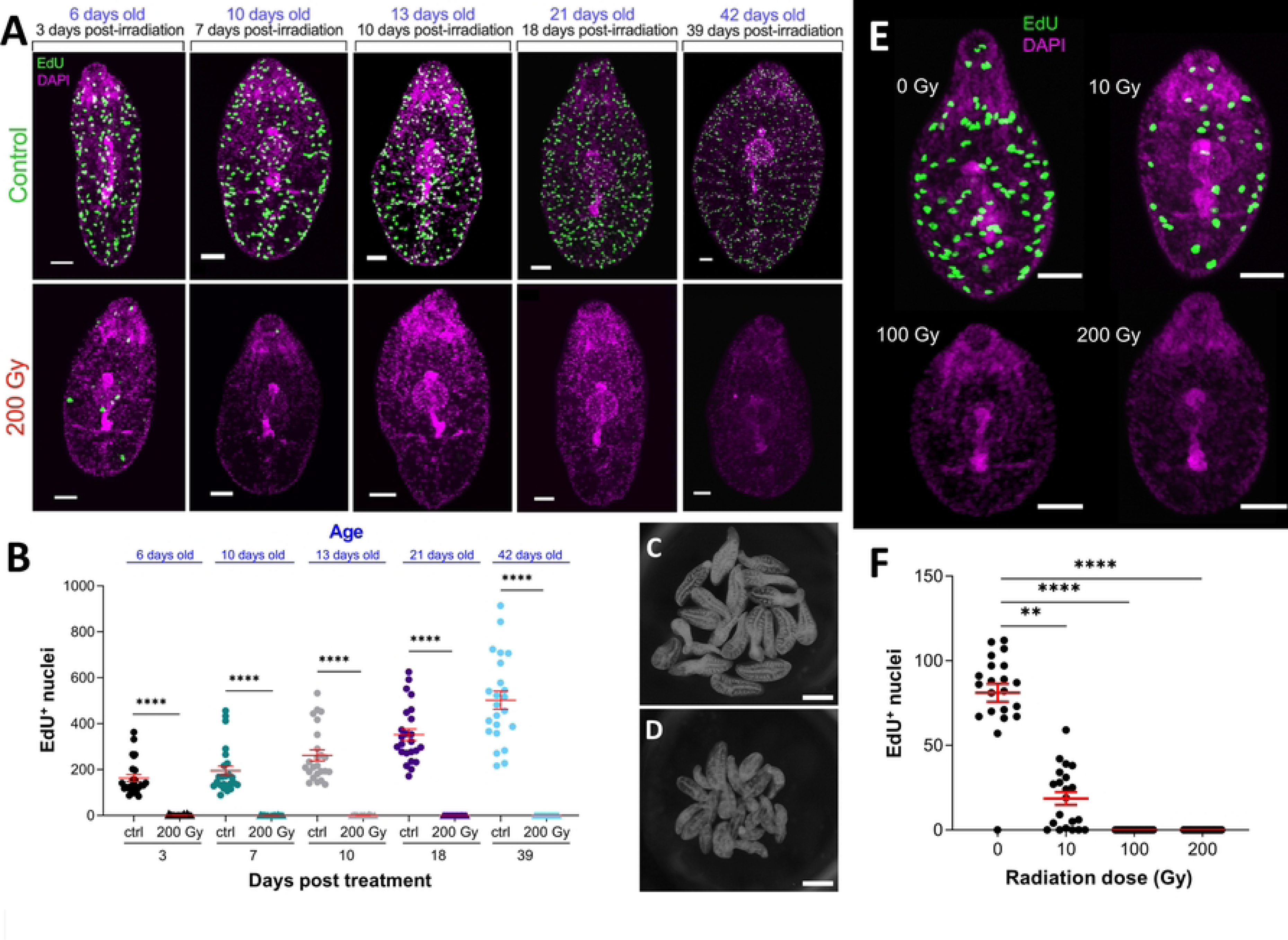
*F. hepatica* neoblast-like cells are inherently linked to juvenile growth. (A) Confocal images of juvenile *F. hepatica* cultured for nine days in either growth inducing media (+CS) or non-growth inducing media (-CS) with labelling of proliferative cells (EdU, green) and all nuclei (DAPI, magenta). (B) Comparison of # EdU^+^ nuclei in juveniles maintained in +CS and -CS shows that growth inducing +CS leads to a significant increase in the number of EdU^+^ nuclei (unpaired t test), scale bars = 50 µm. (C) Plot of # EdU^+^ cells against worm area (each point represents an individual worm) shows a positive correlation (linear regression analysis, p<0.0001), symbols represent different worms of different ages. ****, p<0.0001.

After seven days of culture, the number of neoblast-like cells steadies (Fig 1B). However, immediately preceding this is a marked jump in the number of EdU^+^ nuclei in growing juveniles (Fig 1A; #EdU^+^ nuclei: +CS 4 days, 60±5; +CS 7 days, 279±21). This sudden increase coincides with the timescale for penetration of the liver capsule by *in vivo* juveniles [39]. While direct comparisons between *in vitro* and *in vivo* results are difficult, early studies on the life cycle of *F. hepatica in vivo* found that there was little growth over the first week following infection [39], with a subsequent acceleration in growth around nine days post infection (a few days after liver capsule penetration). Curiously, this increased proliferative rate coincided with a change in EdU^+^ nuclei morphology where cells of older juveniles appeared smaller. To further examine this the areas of 30 EdU^+^ and EdU^-^ nuclei located in the anterior, ventral sucker and posterior regions were recorded. We found that EdU^+^ nuclei nearly halved in size between three- and ten-days post excystment (S1 Fig, EdU^+^ nuclei area: 3 days, 50±0.7 µm; 10 days, 28±0.3 µm), while over the same period there was a marginal increase in the size of EdU^-^ nuclei (EdU^-^ nuclei area: 3 days, 15±0.3 µm; 10 days, 17±0.3 µm). This suggests that subpopulations of *F. hepatica* neoblast-like cells exist. Heterogenous populations of neoblasts, such as zeta, sigma, nu and gamma [14,43,44], are known to exist in planarians and *S. mansoni* neoblasts are also known to be heterogenous [27–29]. Indeed, Hayashi et al. [18] identified two different neoblast sizes and hypothesised that the smaller X2 gate cells were lineage defined progenitors. If this holds true for *F. hepatica* neoblast-like cells, it suggests the larger, early cells have a greater potency, and that these give rise to the smaller, lineage defined cells observed after around seven days. This is reminiscent of the small group of neoblasts present in young *S. mansoni* schistosomula that serve as the parent cells for all future neoblasts [29].

The EdU staining pattern in +CS worms became more regular as they grew. In younger worms (two days old, Fig 1A) EdU^+^ cells were found throughout the somatic tissue. As worms grew the EdU^+^ cells became more evident on the midline and laterally (Fig 1A). This midline/lateral edge staining pattern is akin to the expression pattern of neoblasts in *S. mediterranea* [10], *Dugesia japonica* [45] and *S. mansoni* juveniles [29]. The few EdU^+^ cells seen in non-growing worms were found sporadically throughout the parenchyma of the worm. A final point to note is that we found ∼3x the number of EdU^+^ nuclei after seven days *in vitro* than we observed in our original publication on *F. hepatica* neoblast-like cells [35]. We believe this is due to differences in the length of time that juveniles were incubated in EdU. Here, juveniles were only incubated in EdU for 24 h, whereas previously juveniles were incubated in CS50 with EdU continuously for seven days [35]. Thymidine analogues have been shown in mammalian systems to disrupt the cell cycle and neurogenesis, as well as reducing cell viability [46–48].

### *F. hepatica* neoblast-like cells demonstrate traits associated with differentiation

Differentiation/histogenesis is a key feature of stem cells, since through asymmetric division some daughter cells go on to differentiate into new tissue types. We explored whether the neoblast-like cells of *F. hepatica* were capable of differentiation by carrying out pulse-chase experiments where juveniles were incubated in EdU for 24 h, and then returned to CS50 before fixation/imaging several days later. Fig 2A shows how neoblast-like cells in a three-day old juvenile were found throughout the worm parenchyma. However, a ‘chase’ (EdU exposure, washout and subsequent further culture) interval across a week showed these cells migrating and forming a distinct pattern along the worm’s anterior-posterior midline and lateral edges. Our previous experience with *in vitro* cultured worms revealed that they do not readily develop reproductive structures, but that they do develop extensive gut tissue as well as altering their tegument morphology [35]. Studies in *S. mansoni* have shown that neoblasts contribute to gut development, with ∼40% neoblasts also destined for the tegument [27,28,30]. Considering the midline and lateral accumulation of EdU^+^ cells after a week, we hypothesise that the migrating neoblast-like cells seen here in *F. hepatica* are also contributing to gut/tegument development. Curiously, the ‘chase’ pattern of neoblast-like cells along the worm midline and lateral edges is very similar to the staining pattern in juveniles ≥7 days old immediately after EdU incubation (Fig 1A). This suggests that the neoblast-like cells present in young juveniles (1-3 days old) may migrate to the anterior-posterior midline/lateral edges, and from there act as multipotent stem cells that are partially lineage defined.

**Fig 2.**
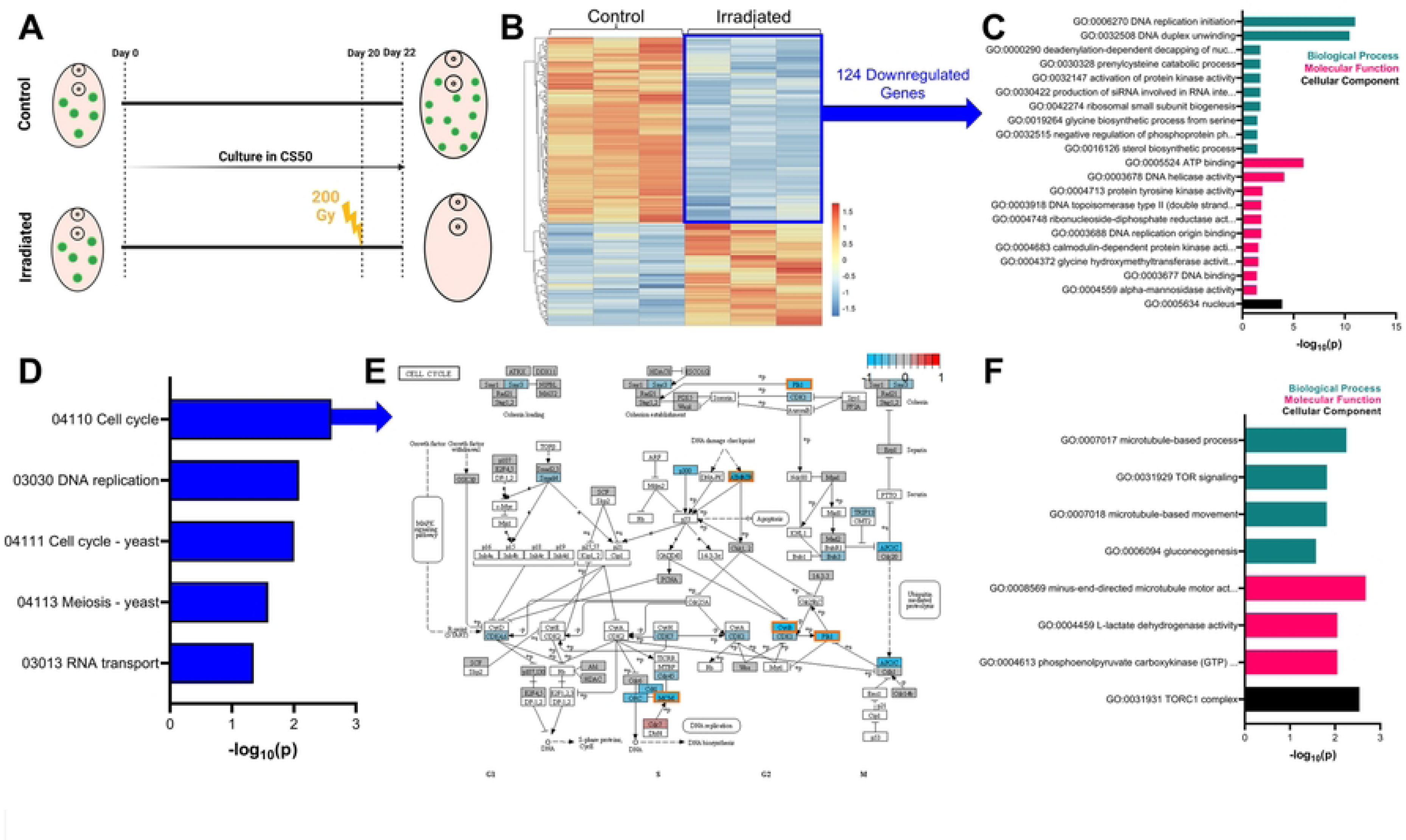
Evidence of differentiation by *F. hepatica* neoblast-like cells. (A) Confocal images with EdU labelling (green) show pulse-chase trial where EdU^+^ cells migrate to worm midline and lateral margins over a one-week chase period (blue = # of days post-pulse). (B) The dissociation of juveniles just after EdU incubation illustrates how EdU^+^ cells (images on left) have ovoid morphology with scant cytoplasm, in contrast to EdU^-^ cells (image on right). (C) The dissociation of juveniles 72 hours after EdU incubation illustrates how EdU^+^ cells (images on left) have a similar morphology to EdU^-^ cells (image on right), suggesting cell differentiation. Differential interference contrast microscopy and DAPI counterstain (magenta) used to provide context of images. Scale bars; A = 50 µm; B/C = 10 µm.

We currently lack tissue specific labels for *F. hepatica* and so to further explore the differentiation potential of the neoblast-like cells we examined the morphology of EdU^+^ cells before and after ‘chase’ periods. Fig 2B shows how EdU^+^ cells fixed and stained immediately following EdU incubation have a large nucleus with scant cytoplasm and are smaller than EdU^-^ cells, both known neoblast features [9,25,49]. However, following a three-day ‘chase’ EdU^+^ cells appeared to be larger and to possess a greater cytoplasm volume, akin to the EdU^-^ cells (Fig 2C), indicating that the neoblast-like cells of juvenile *F. hepatica* are growing, and likely differentiating into new cell types.

### Dual-labelling suggests *F. hepatica* neoblast-like cells are capable of self-renewal

Stem cells must be capable of self-renewal. To test this in *F. hepatica* we used two thymidine analogues at different time points to assess if neoblast-like cells were capable of replication days apart. Three-week-old juveniles were incubated in EdU for six hours, followed by a 48-hour chase before a 24-hour incubation in BrdU, a second thymidine analogue (Fig 3A). This chase period was chosen to give cells time to complete mitosis before undergoing the cell cycle once again. Fig 3B shows fluke with numerous nuclei positive for both EdU^+^ and BrdU^+^ (Fig 3E; mean percentage of EdU^+^/BrdU^+^ nuclei: 53%±2.3%). In *S. mansoni* adults ∼41% of cells were EdU^+^/BrdU^+^ when using a similar staining protocol [25], suggesting a similar self-replication rate in both species. We also recorded the presence of pairs of cells with differing staining combinations (Fig 3C & D). These pairs of cells are like the ‘doublets’ identified by Collins et al. [25], and their presence supports evidence of the mitotic nature of EdU^+^ cells in *F. hepatica*. Asymmetric division of neoblasts is crucial for maintenance of both the stem cell line and differentiating cells [50]. Fig 3D demonstrates an EdU^+^ doublet pairing of EdU^+^/BrdU^+^ and EdU^+^/BrdU^-^, which indicates asymmetric division of cells where one is for self-replication, while the other is destined for differentiation. We also noted that the percentage of proliferating cells incorporating EdU alone was similar to that for BrdU alone, despite an 18-hour difference in incubation times (Fig 3E; proliferating cells: EdU^+^ alone, 21±3.3%; BrdU^+^ alone, 26±2.2%). This could be due to either: (i) EdU^+^ cells cycling numerous times during the intervening 64-hour chase, leading to staining of their daughter cells, or (ii) the thymidine analogues limiting the rate of proliferation [46,47].

**Fig 3.**
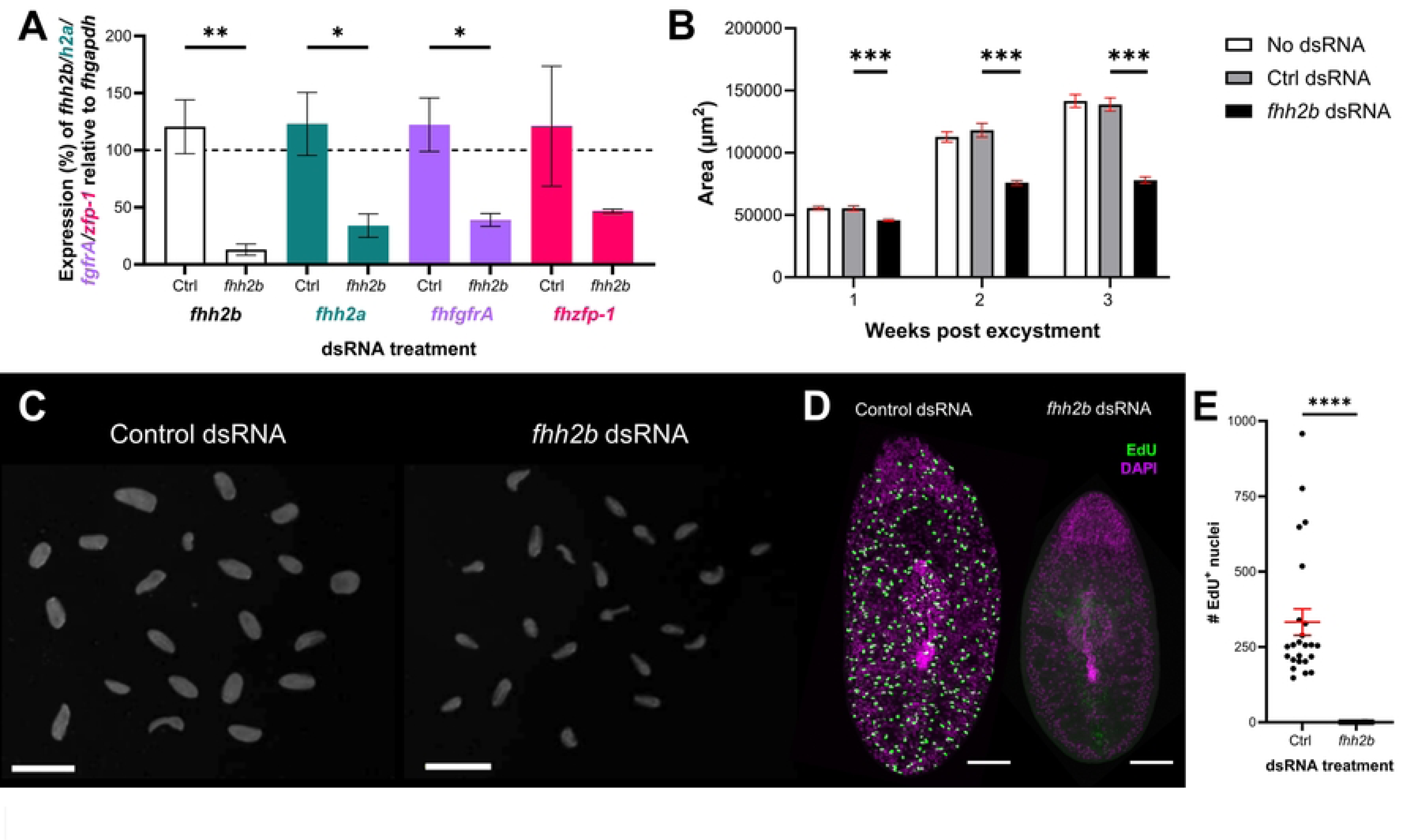
*F. hepatica* neoblast-like cells are capable of self-renewal. (A) Timeline of dual labelling with the thymidine analogues EdU and BrdU where colocalisation would be indicative of self-renewal. (B) Confocal images of EdU (green), BrdU (red) and merged channels showing colocalisation of EdU/BrdU (yellow/orange) in worms treated according to above timeline, supporting the presence of self-renewing cells. Scale bar = 100 µm. (C) Confocal image of the mid-region of a juvenile labelled with EdU/BrdU according to the above timeline shows colocalisation of EdU/BrdU (orange arrowheads) as well as the presence of doublets (white arrowheads). Scale bar = 10 µm. (D) Asymmetric pairings of EdU^+^/EdU^+^ and BrdU^-^/BrdU^+^ cells (white arrowhead). Scale bar = 10 µm. (E) Based on the percentage of proliferating cells that were EdU^+^, BrdU^+^ or EdU/BrdU^+^, it was evident that ∼50% of these cells had self-replicated within the experimental timeframe.

### Ionising radiation ablates *F. hepatica* neoblast-like cells, disrupting *in vitro* growth

The mitotic activity of stem cells makes them particularly susceptible to damage through irradiation with the neoblasts of planarians and parasitic worms amenable to ablation following irradiation [25,33,51,52]. *F. hepatica* neoblast-like cells appeared to be similarly ablated following irradiation, with doses of 100-200 Gy preventing neoblast-like cell proliferation (S2 Fig) but having no effect on juvenile survival in the 4 weeks following irradiation. While other parasitic flatworms have been reported to survive radiation doses of 200 Gy [25,33], planarians are killed by ∼60 Gy [16], and rats by doses <20 Gy [53]. Why parasitic flatworms have evolved to resist the damaging effects of such high levels of radiation remains a mystery, though *F. hepatica* is known for resisting free radical damage [54], which may be linked to their defence against host immune system leukocytes [55]. We also found that if worms were incubated with EdU prior to irradiation this did not completely ablate the EdU^+^ cell population, though it was significantly reduced (S3 Fig). This reduction in the number of EdU^+^ cells may be due to the ablation of neoblast-like cell proliferation, while the remaining EdU^+^ cells were those daughter cells that were now differentiating.

We examined whether these neoblast-like cells recovered in the weeks following irradiation. Three-day old juveniles were subjected to radiation and then incubated in EdU and fixed at time points over the subsequent 39 days. This showed that neoblast-like cell proliferation did not recover following ablation within this time period (Fig 4A & B). Though worms remained viable, they appeared to be increasingly compromised over the weeks following irradiation with reduced growth/development (Fig 4C & D). This disruption of growth is likely due to a lack of neoblast-like cells available for differentiation into new tissues. In planarians, the localised ablation of neoblasts results in necrosis in the afflicted area, though amputation encourages the regeneration and reestablishment of neoblast populations [56]. Extended culture of irradiated *F. hepatica* may reveal if radiation ultimately leads to death.

**Fig 4.**
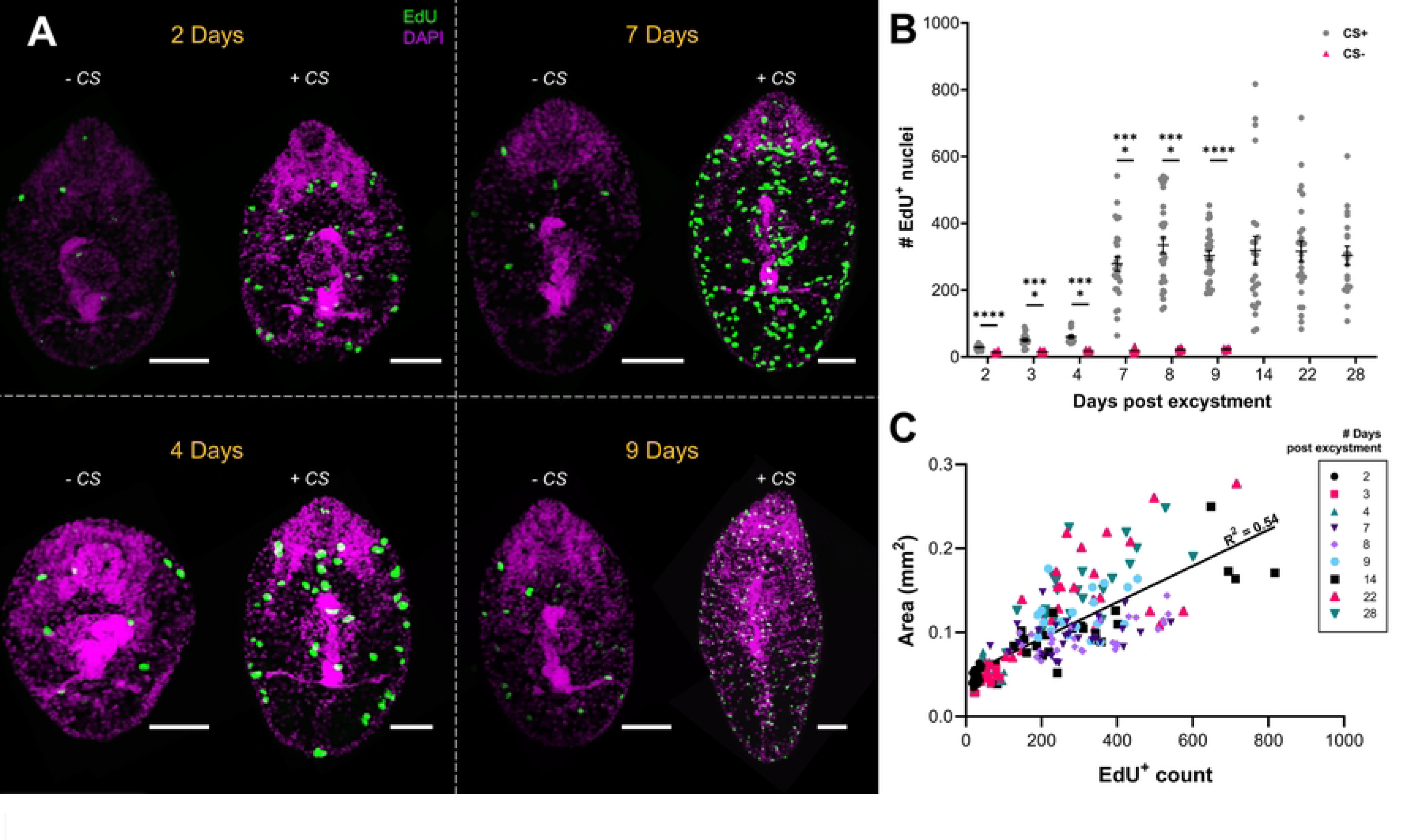
The neoblast-like cells of *F. hepatica* are sensitive to ionising radiation. (A) Confocal images of three-day old juvenile *F. hepatica* irradiated with 200 Gy before 39 further days of culture shows that irradiated worms do not recover EdU^+^ nuclei (blue text = age; black text = # days post irradiation). EdU (green), DAPI (magenta); Scale bars = 50 µm. (B) # EdU^+^ nuclei in 200 Gy irradiated and non-irradiated worms tracked over 39 days following irradiation (blue text = age; black text = #days post irradiation) shows significant reduction in # EdU^+^ at all time points (Kruskal-Wallis and Dunn’s posthoc test). (C/D) Dark-field images of control (C) and 200 Gy irradiated (D) worms 39 days post irradiation shows reduced growth in irradiated worms; scale bars = 200 µm. (E) Confocal images of two-day-old juvenile *F. hepatica* treated with 0 – 200 Gy radiation as metacercariae prior to excystment demonstrates dose dependent effects of irradiation on EdU^+^ nuclei (green), DAPI (magenta); scale bars = 50 µm. (F) # EdU^+^ nuclei in two-day-old juvenile *F. hepatica* treated with 10, 100 or 200 Gy radiation as metacercariae (Kruskal-Wallis and Dunn’s posthoc test). **, p<0.01; ****, p<0.0001.

Unlike *S. mansoni*, *F. hepatica* have a dormant infective life stage, the metacercaria, that follows the cercarial stage. This afforded us the opportunity to examine if proliferating cells were present within this dormant life stage by exposing metacercariae to radiation prior to excystment. Fig 4E shows how doses of 10-200 Gy prior to excystment reduce neoblast-like cell numbers, with even a low dosage (10 Gy) significantly depleting numbers (Fig 4F). While studies on the metacercarial stage are limited, previous work has identified transcripts in metacercariae [57]. Our evidence of proliferation suggests that, despite being dormant, metacercariae are transcriptionally active and are primed to increase proliferation following excystment and host invasion.

Irradiation of *F. hepatica* metacercaria has previously been trialled in attempts to develop a vaccine [58–60]. While these attempts were unsuccessful in developing viable vaccines, they provided insights into the effects of radiation on worms. Irradiation of metacercariae prior to infection resulted in increased host immune responses and slowed worm growth/development [58,60]. In fact, radiation doses >100 Gy resulted in <1% of *F. hepatica* reaching maturity [60], and 30-40 Gy significantly disrupted tegument development [59]. Considering these observations with our own data, the disruption of neoblast-like cells of metacercariae via irradiation is likely to have contributed to the limited development of fluke *in vivo*. If neoblast-like cells are as crucial for tegument maintenance/development as they are in *S. mansoni* [27,28], then imbalance in this system would have serious implications for the host-parasite relationship, potentially allowing the host immune system to identify and target migrating juveniles.

### Irradiated worms downregulate key stem cell associated genes

To further explore the biology of the neoblast-like cells we subjected juvenile *F. hepatica* to transcriptomics following radiation. Worms were grown for three weeks *in vitro* and then irradiated with 200 Gy prior to RNA extraction and sequencing 48 hours later (Fig 5A). We found 124 genes were downregulated following irradiation when compared to non-irradiated, time-matched controls (Fig 5B; DESeq2, p<0.05). We hypothesised that these downregulated transcripts were associated with neoblast-like cells that appear to be ablated following irradiation. TOPGO analysis showed that genes involved in nucleotide interactions, especially those involved in the cell cycle and located in the nucleus were over-represented among downregulated genes (Fig 5C). We utilised BLAST databases and our own manual annotations to identify the downregulated genes (S2 Table). Two fibroblast growth factor receptors (*fgfrA* and *fgfrB*) were identified among the top 40 downregulated transcripts. FGFRs are receptor tyrosine kinases for fibroblast growth factors (FGFs) and their signalling pathway in mammals plays crucial roles in tissue homeostasis, endocrine function, and wound repair [61]. Their dysregulation can lead to cancer, and so they are considered a favourable therapeutic target for chemotherapy [61,62]. The FGFR expressing cells of planarians are irradiation sensitive [63], and direct brain development [64–66]. *E. granulosus* do not appear to express endogenous FGFs and may use host FGFs as ligands for their FGFRs to drive growth/development via their stem cells [67]. A homologue of *F. hepatica fgfrA* was downregulated following irradiation in *S. mansoni* [25], and was found to be associated with their neoblasts both through expression and RNAi studies [24,25]. Curiously, while *fhfgfrA* and *fgfrB* were downregulated, we noted that a third *fgfr* (*fhfgfrC*) was upregulated following irradiation. Further studies into these *fgfrs* may show if differing responses to irradiation are linked to different functions/expression patterns within the worms. Another proliferation associated kinase, polo-like kinase 1 (*plk1*), was also downregulated following irradiation (S2 Table). PLK1, a serine/threonine kinase, is crucial to the cell cycle, playing roles in the regulation of mitosis onset, spindle assembly and cytokinesis [68]. The dysregulation of PLKs is again linked to the over proliferation of cancer cells and makes it an appealing target for novel drugs [69]. A recent study in *F. hepatica* demonstrated how a PLK inhibitor reduced movement in juvenile fluke and egg production in adult worms [70]; coincidentally, *S. mansoni* PLKs are known to be associated with germline cells [71,72]. In total we identified 10 downregulated kinases following irradiation. With a plethora of kinase inhibitors being developed for cancer treatments and other diseases associated with over proliferation (e.g. idiopathic pulmonary fibrosis) [73], it is possible that some could be repurposed as anthelmintics to disrupt parasitic helminth growth/development within the host.

**Fig 5.**
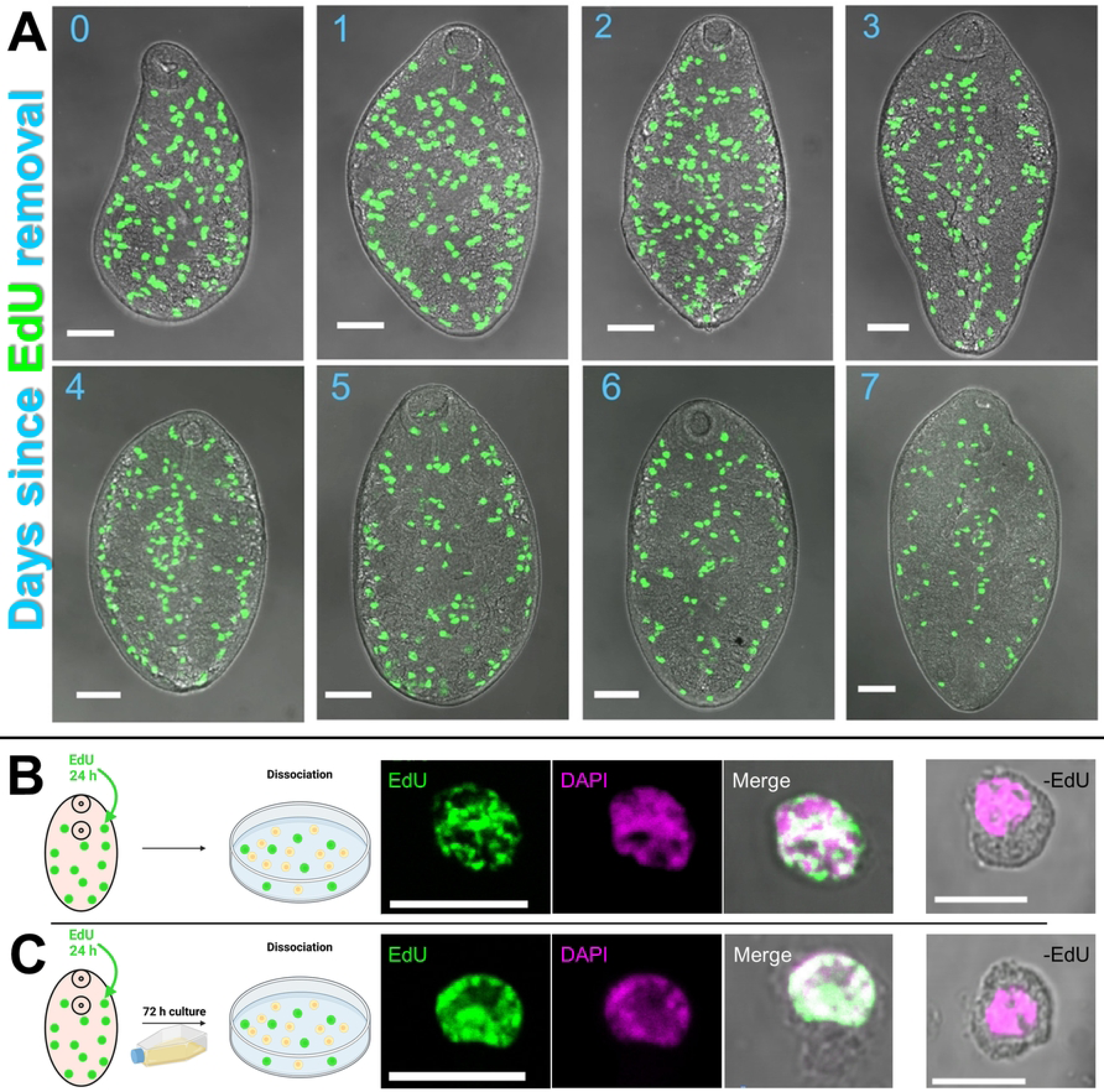
Irradiation of juvenile *F. hepatica* downregulates stem cell associated transcripts. (A) Timeline of experiment to generate irradiated transcriptomes where juvenile *F. hepatica* were grown for 20 days *in vitro* before treatment group was irradiated with 200 Gys to ablate stem cells. (B) Heatmap of significantly differentially expressed genes, as identified by DESeq2, highlights 124 downregulated genes following irradiation. (C) TOPGO analysis shows genes associated with cell replication, and localised to nucleus, overrepresented among downregulated genes following irradiation. (D) Significantly downregulated KEGG pathways following irradiation. (E) KEGG cell cycle pathway shows that many pathway components are downregulated (upregulated, red; no change, grey; downregulated, blue; unassigned KEGG ID, white; differentially expressed, orange box). (F) TOPGO analysis shows genes associated with microtubules and TOR signalling overrepresented among upregulated genes following irradiation.

A second group of genes that were significantly downregulated were transcription factors. Transcription factors have not classically been regarded as drug targets due to difficulties in the development of small molecules against targets with broad and featureless interaction sites [74]. However, the specificity of transcription factors makes them potentially more appealing as drug targets than upstream receptors (e.g. kinases) [75]. We found at least 12 transcription factors downregulated following irradiation (S2 Table), including a kruppel-like factor (*klf*), *hesl*, *sox*, and several zinc-finger proteins (*zfp*). While the transcriptional targets of these proteins in *F. hepatica* are unknown, a number have been found to contribute to development in planarians [12,14,76], and *zfp-1* and *zfp1-1* are associated with the development of epithelial cells in *S. mansoni* [28], while *foxA* contributes to oesophageal gland development [31]. This highlights the potential for the occurrence of putative drug targets among the transcription factors regulating juvenile growth/development. One additional feature of our downregulated genes is that 28% remained unidentified (unknown with no BLAST hit <0.001 against NCBI landmark and swissprot databases). While these genes remain the most elusive, and some may well be mispredictions, some could also represent attractive targets, given that they appear to be *Fasciola* or parasitic flatworm specific. They could be involved in important biological processes, such as host-parasite interactions, and warrant further investigation given their lack of homology to genes/proteins in host organisms.

We also wanted to compare our list of significantly downregulated genes against those downregulated following irradiation in *S. mansoni* [25]. Although comparisons between transcriptomic datasets is imperfect, compounded by the fact that *S. mansoni* work was carried out in *ex vivo* adults while we used *in vitro* juvenile *F. hepatica*, we found 27 genes downregulated in both datasets (S3 Table). Among those genes downregulated in both the schistosome and *Fasciola* studies were *fgfrA*, *plk*, *p53* and members of the mini-chromosome maintenance complex. This alignment encourages confidence that the downregulated genes in our transcriptomic dataset are representative of transcripts associated with *F. hepatica* neoblast-like cells and provides a list of core, conserved parasitic flatworm stem cell transcripts. However, we noted that certain neoblast associated genes we expected to find downregulated in our datasets, such as *nanos2*, were absent. Published studies showed that *nanos2* was differentially expressed in more mitotically active *F. hepatica* [38] and was downregulated in *S. mansoni* following irradiation [25]. Upon investigation of transcript per million data (TPM; S4 Table) we found that *nanos2* absence was linked to low expression across both treatment groups, likely due to the limited proliferation rate in our small *in vitro* worms (*fhnanos2* TPM: control, 5.6±0.2; irradiated, 3.1±0.4). Therefore, we identified the *F. hepatica* homologues of *S. mansoni* radiation-affected genes [25] and examined their expression profiles in our dataset (S4 Fig). We found ∼50% of the *S. mansoni* homologues exhibited marked downregulation in expression following irradiation. Finally, we identified the *S. mansoni* homologues of all 124 *F. hepatica* downregulated genes and examined *S. mansoni* single cell datasets [30,77] to ascertain if transcripts were neoblast associated or not (S4 Fig), finding that 68 genes appear to be neoblast-linked in schistosomes. Together, these data support our hypothesis that the irradiation-sensitive cells of *F. hepatica* are comparable to the neoblasts reported in other flatworm species.

We confirmed our DESeq2 findings by examining which KEGG pathways were significantly up/downregulated by irradiation through mapping KEGG IDs onto *F. hepatica* predicted proteins. We found that KEGG pathways associated with the cell cycle were downregulated (Fig 5D), in particular the cell cycle itself (Fig 5E) and DNA replication pathways. This highlighted that important cell cycle components, such as cyclin-dependent kinase 1 (*cdk1*), were downregulated following irradiation, even though they were not classed as significantly downregulated by DESeq2. This second method of analysis gave further verification of results showing that the most mitotically active cells, the neoblast-like cells, were those ablated following irradiation and so downregulated transcripts are likely to be markers of neoblast-like cells.

We also examined upregulated genes/pathways following irradiation. No KEGG pathways were found to be upregulated, but we found 69 upregulated genes, though 49% of these remain unidentified due to a lack of BLAST hits against model organism databases. While there was a limited pool of upregulated genes, TOPGO analysis suggested that GO terms associated with microtubules and the TORC signalling were upregulated. Upregulation of TORC signalling was unexpected, as this pathway typically regulates stem cell proliferation in response to nutrient availability [78]. However, the TORC pathway is involved in blastema formation in planarians following wounding [79,80], and so could be involved in repairing damage done via irradiation in *F. hepatica*.

### RNAi-mediated knockdown of *h2b* ablates *F. hepatica* stem cells and limits growth

Irradiation is an extremely useful tool for the study of stem cells/neoblasts, but it is somewhat of a blunt instrument as it can also damage differentiated cells. Therefore, we used RNAi to silence histone *h2b*, an important cell cycle component due to its role in transcription, as an alternative method of disrupting the *F. hepatica* neoblast-like cells [17]. Fig 6A shows how dsRNA targeting *fhh2b* resulted not only in significant reduction of the *fhh2b* transcript, but also of histone *h2a* and *fhfgfrA* transcripts. Additionally, we observed a slight reduction in the expression of *fhzfp-1*, a homologue of the *zfp* in *S. mansoni* that contributes to epidermal cell lineages [28]. Silencing of *fhh2b* also resulted in a marked reduction in fluke growth (Fig 6B & C) just one week after commencement of the trial, followed by very limited growth over the subsequent two-week period. We hypothesised that this was related to the disruption of neoblast-like stem cell proliferation, confirmed by the absence of EdU^+^ cells in *fhh2b* dsRNA treated worms (Fig 6D & E). These results suggest that the knockdown of a key cell cycle regulator has similar impacts to that of irradiation, as has been seen previously in both free living and parasitic flatworms [17,25,33]. Again, we found that *fhfgfrA* expression was reduced following the disruption of neoblast-like cells, which demonstrates how closely linked this gene is to growth and development in *F. hepatica*. Indeed, in *S. mansoni* this gene was expressed in 99% of all neoblasts [25], and as such may be a gene worth considering for its drug target candidature. The severe disruption to growth and development seen here through precise targeting of a neoblast-like cell regulator also illustrates a proof of concept that these cells should be considered as viable targets for novel flukicides.

**Fig 6.**
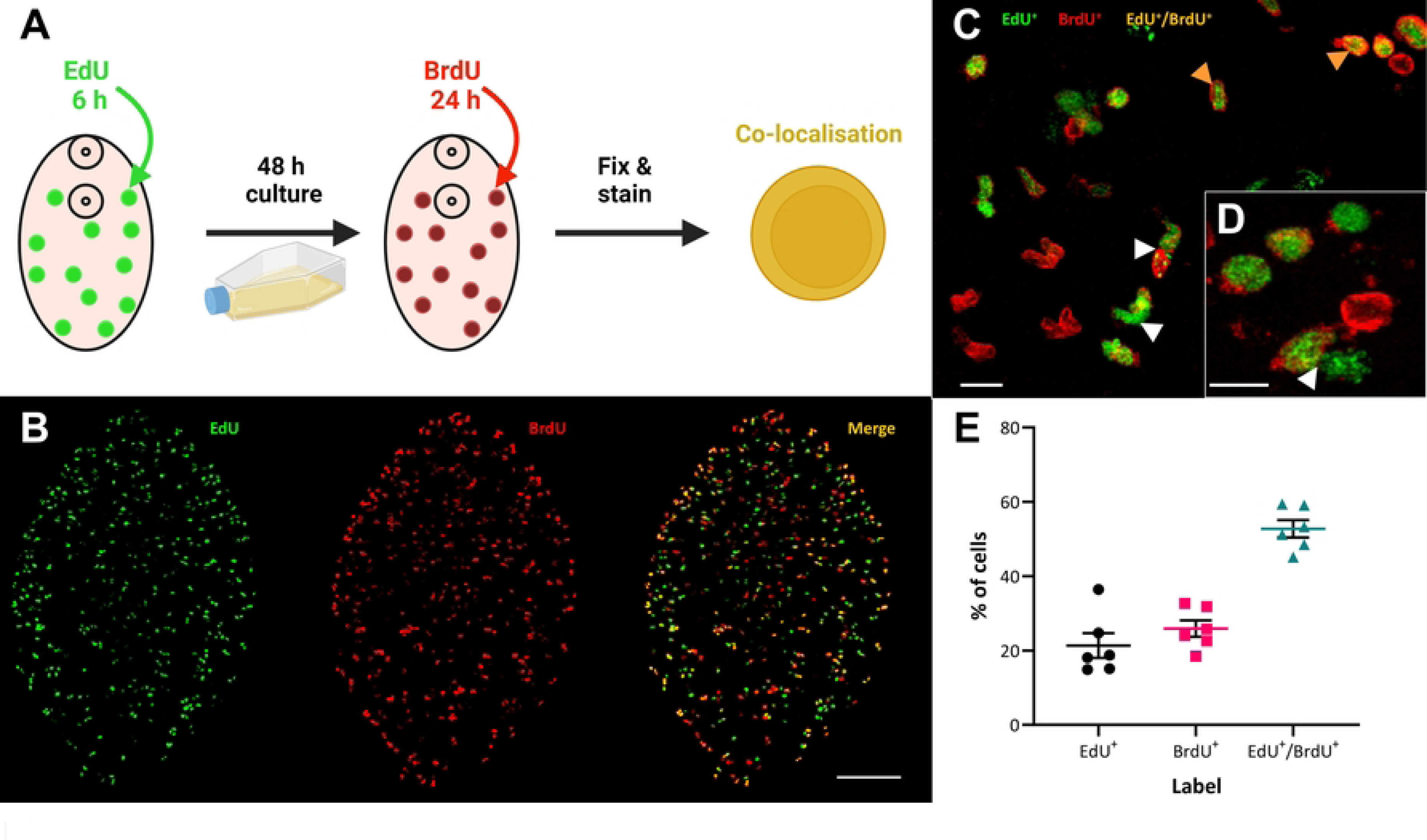
RNAi mediated knockdown of *fhh2b* significantly reduces stem cell associated transcripts and neoblast-like nuclei in juvenile *F. hepatica in vitro*. (A) Expression of *fhh2b* (white), *fhh2a* (green), *fhfgfrA* (purple) and *fhzfp-1* (pink) relative to *fhgapdh* in *in vitro* juvenile *F. hepatica* following three weeks of repeated exposures to *fhh2b* dsRNA shows significant knockdown of stem cell associated transcripts *fhh2b*, *fhh2a* and *fhfgfrA* relative to control dsRNA treated worms (unpaired t tests). (B) The *in vitro* growth (area in µm^2^) of juvenile *F. hepatica* was significantly reduced after three weeks of repeated exposure to *fhh2b* dsRNA (Kruskal-Wallis and Dunn’s posthoc tests). (C) Darkfield images of three-week old *in vitro* juvenile *F. hepatica* show how juveniles treated repeatedly with *fhh2b* dsRNA are significantly less developed than those treated with control dsRNA; scale bars = 1 mm. (D) Confocal images of EdU staining (green) in three-week-old *in vitro F. hepatica* treated with *fhhh2b* dsRNA show ablation of EdU^+^ nuclei compared to control dsRNA treated worms; scale bars = 100 µm, DAPI counterstain (magenta). (E) # EdU^+^ nuclei are significantly reduced in three-week-old *F. hepatica* juveniles following repeated treatment with *fhh2b* dsRNA (Mann-Whitney U test). *, p<0.05; **, p<0.01; ***, p<0.001; ****, p<0.0001.

## Conclusion

The results throughout this paper clearly demonstrate the key characteristics of the somatic neoblast-like cells found in *F. hepatica*. They are (i) capable of differentiation, (ii) capable of self-renewal, and (iii) sensitive to ionising radiation, all defining characteristics of stem cells. However, it is their connection to the somatic growth and development of the invasive, and highly damaging, juvenile *F. hepatica* that makes them attractive resources for novel flukicide targets. Here, we have identified several sets of genes that represent putative drug targets, including several kinases clearly linked to the stem cells of several parasitic flatworms. Repurposing anthelmintics for cancer treatment is now a growing field of research [81], but it is similarly appealing to consider the repurposing of anti-cancer drugs (and compounds from anticancer drug development campaigns) as anthelmintics, with kinases having been previously touted as potential anthelmintic targets [82]. This study has scrutinised the fundamental biology of *F. hepatica* somatic neoblast-like stem cells, underpinning our understanding of their pathogen biology and supporting efforts to develop novel therapeutics for a globally import parasite.

## Materials and Methods

### *F. hepatica* excystment and *in vitro* culture

Italian strain *F. hepatica* (Ridgeway Research, UK) were excysted as described by McVeigh *et al.* and McCusker *et al.* [83,84]. A full excystment protocol is available at http://dx.doi.org/10.17504/protocols.io.14egn212qg5d/v1. Briefly, metacercariae had their outer wall mechanically removed prior to 2-3 min bleaching in 10% sodium hypochlorite, which was washed off with >3x rinses in reverse osmosis (RO) water. Cysts, with only their inner wall remaining, were incubated in excystment solution for one hour. Excysting newly excysted juveniles (NEJs) were washed in RPMI (#11835105, Thermo Fisher Scientific) prior to culture in 96 well plates in CS50 (50% chicken serum (#16110082, Thermo Fisher Scientific), 50% RPMI with antibiotic/antimycotic (#A5955, Sigma Aldrich) as reported previously [35]. Throughout trials culture media was changed three times/week. NEJ is used to refer to worms within 24 hours of excystment and juvenile for worms cultured beyond this point.

### 5-ethynyl-2’-deoxyuridine (EdU) Labelling

Juveniles were incubated in 500 µM EdU (10 mM EdU stock resuspended in PBS) for 24 hours (final dilution in either CS50 or RPMI depending on the trial), after which point worms were either fixed immediately, or returned to CS50 for further culture. Juveniles ≤1-week old were free-fixed in 4% PFA (4% paraformaldehyde in PBS) for four hours at RT. Juveniles >1-week old were flat-fixed under a coverslip for 10 min in 4% PFA prior to four hours free-fixing at RT. Following fixation, juveniles were permeabilised in PBST (0.5% Triton X-100 in PBS) for 30 min at RT before EdU detection, using the reaction cocktail from the Click-iT EdU Cell Proliferation Kit with an Alexa Fluor 488 dye (#C10337, Thermo Fisher Scientific) for 30 min at RT. Worms were then washed twice in PBST before incubation in 5 µg/mL 4′,6-diamidino-2-phenylindole (DAPI) in PBS for 20 min to label nuclear DNA. Worms were mounted in Vectashield (Vector Laboratories) and imaged using a Leica TCS SP8 inverted microscope. Maximum z stack projections were exported for analysis.

For pulse-chase experiments worms were treated with EdU for 24 h, after which EdU-containing media was rinsed off using three 1 mL RPMI washes. Worms were then returned to culture and fixed at desired timepoint for staining.

### Cell dissociation and staining

Thirty three-week-old juveniles were incubated in EdU for 24 hours as stated previously. Following EdU exposure juveniles were washed 3x in 1 mL RPMI to remove chicken serum before being cut into three segments using a sterile scalpel. Segments were dissociated in hydrophobic tubes containing 200 µL of 2.5 mg/mL Liberase DH (#5401054001, Sigma Aldrich) while being shaken at 300 rpm for 1 hour 30 min at 37°C, with brief vortexing every 30 min. The resulting cell suspension was filtered through a 70 µm pore Flowmi cell strainer before centrifugation at x300 *g* for 5 min at 4°C. The supernatant was removed, and the cell pellet resuspended in 100 µL of 4% PFA, then left to fix for 30 min at RT.

Fixed cells were spotted onto Superfrost Plus Adhesion Microscope Slides (Thermo Fisher Scientific) and air dried for 5 min at RT. Cells were permeabilised for 30 min in 200 µL of 0.3% Triton X-100 in PBS (PBSTx). PBSTx was rinsed off with 2x washes in 200 µL PBS prior to EdU detection and DAPI counterstaining (carried out as stated above). Vectashield was applied and cells were sealed with a coverslip prior to imaging as stated previously.

### EdU/ Bromodeoxyuridine (BrdU) colocalisation

Three-week-old juvenile *F. hepatica* were incubated in EdU for 6 hours (the ‘pulse’) as previously described, before being returned to culture in CS50. After 48 hours (the ‘chase’) juveniles were exposed to 500 µM BrdU (10 mM BrdU stock resuspended in PBS) in CS50 for 24 h, before fixation in 4% PFA as previously described. Specimens were permeabilised with PBST for 30 min at RT, followed by a 15 min incubation in proteinase K (0.1% SDS and 11 µg/mL Proteinase K in PBST) at RT. Proteinase K was removed, and juveniles were post-fixed in 4% PFA for 10 min, prior to 2x 5 min RT washes in PBST. To expose incorporated BrdU for labelling, DNA denaturation was achieved through a 30 min incubation in 2N HCl in PBST at 37°C. HCl was removed with 4x 10 min PBS washes at RT. EdU detection was achieved using an adapted version of the *S. mansoni* EdU staining protocol [85], available at https://dx.doi.org/10.17504/protocols.io.eq2lyjnrrlx9/v1, followed by two 5 min washes in BSA (0.3% bovine serum albumin in PBS). BrdU was immunolocalised through incubation in an anti-BrdU monoclonal antibody (1:500 dilution in BSA, MoBU-1, Abcam) for 72 hours at 4°C. Worms were then washed three times at RT in PBST to remove primary antibody, before incubation in goat anti-mouse IgG conjugated to Alexa Fluor 647 (Life Technologies) at 1:100 (diluted in BSA) for 48 hours at 4°C. After three further PBST washes and DAPI counterstaining worms were mounted and imaged as previously described.

Z stacks were imported into IMARIS software (v9.9) to quantify the number of EdU^+^ and BrdU^+^ cells. Briefly, cells were quantified by identifying ‘spots’ of absolute intensity that were 4 µm in diameter (quality, >80; region threshold, 410). Colocalised cells were counted as cells where the spot centre points were within 1 µm of each other.

### Irradiation

Juvenile worms and metacercariae (with outer cyst wall removed) were irradiated in round bottomed 96-well plates in RPMI or ddH_2_O, respectively. A 160 kVp X-ray source with a 0.8 mm beryllium filter (Faxitron CP-160) was used. Worms were irradiated at a height 11 cm from the source, providing a uniform dose rate of 4.5 Gy /min. Total dose administered ranged from 10 Gy to 200 Gy. Post-radiation treatment juveniles were further cultured in CS50 and metacercariae processed for excystment as described above.

### Irradiated transcriptome generation and analysis

Worms were excysted and cultured for 20 days as stated previously. On day 20 worms were separated into 6x replicates of 50 worms. Three replicates were subjected to 200 Gy of radiation as described previously, with the remaining replicates acting as controls. Worms were returned to culture for 48 h. On day 22 juveniles were snap frozen in liquid nitrogen and stored at -80°C. Total RNA was extracted using Trizol reagent (#15596026, Thermo Fisher Scientific). RNA samples were quantified and assessed for purity using an AATI fragment analyser by the Genomics Core Technology Unit at Queen’s University Belfast (QUB GTCU, https://www.qub.ac.uk/sites/core-technology-units/Genomics/). The QUB GTCU generated libraries (3x control and 3x irradiated) using the KAPA mRNA HyperPrep Kit (Roche) prior to paired-end sequencing (2 x 50 bp) on an Illumina Nova seq 6000 SP100 with ∼40M reads/sample. Raw read files are available from the European Nucleotide Archive under accession PRJEB64689.

Adaptor sequences were trimmed using Trimmomatic (v.0.39; [86]) and trimmed reads quality was assessed via fastqc (v.0.11.8). The University of Liverpool *F. hepatica* genome [57] and GFF files (PRJEB25283, WBPS15, https://parasite.wormbase.org/Fasciola_hepatica_prjeb25283/Info/Index/) were downloaded from WormBase ParaSite [87] and used for read alignment with HISAT2 (v.2.1.0; [88]). Stringtie (v.1.3.6; [89]) was used to assemble transcripts. Transcripts from different replicates and reference genome were merged to create a new reference file (excluding isoforms). Gene counts were then quantified by Stringtie (v.1.3.6), exported into a format readable by DESeq2 (v.1.34.0), and filtered to remove non-coding genes and genes with <10 counts across all samples.

Analysis of differentially expressed genes was carried out in R (v.4.2.1) by DESeq2 (v.1.34.0; [90]), using default parameters. A false discovery rate (FDR) p-value threshold of 0.05 was used to identify up/downregulated genes following irradiation. Differentially expressed genes (DEGs) were identified through BLASTp (for predicted proteins) or BLASTx (for ‘novel’ identified genes, listed as MSTRG; S1 File) searches against the NCBI landmark and swissprot databases (p<0.01). Manual annotations of *F. hepatica* genes, both published and in-house, were also used to identify genes. Additionally, DEGs were also used in BLAST searches against the *S. mansoni* genome (v8, PRJEA36577, WBPS15, https://parasite.wormbase.org/Schistosoma_mansoni_prjea36577/Info/Index/) to identify homologues.

GO terms associated with the *F. hepatica* genome assembly were downloaded from WormBase ParaSite (PRJEB25283, WBPS15) to perform TOPGO enrichment analysis (v.2.46) on differentially expressed genes (TOPGO analysis parameters: FDR <0.05, method = weight01, statistic = fisher) within R. Additionally, *F. hepatica* predicted proteins (PRJEB25283, WBPS15) were run through BlastKOALA (https://www.kegg.jp/blastkoala/) to assign predicted KEGG IDs. R programs gage (v.2.44) and pathview (v.1.34) were then used with fold change data from all genes (regardless of differential expression status) to identify up/downregulated KEGG pathways in irradiated worms. Custom R scripts available at https://github.com/pmccusker09/F.hepatica_irradiated_transcriptome_R_analysis.git.

### Histone 2B (*h2b*) RNAi

cDNA for dsRNA templates was obtained from juvenile *F. hepatica* cultured for three weeks in CS50. Worms were snap frozen in liquid nitrogen before being ground into a fine powder for 1 min by a stainless-steel bead in a TissueLyser LT (Qiagen) operating at 50 oscillations per second. mRNA was extracted using the Dynabeads mRNA DIRECT Purification Kit (#61012, Thermo Fisher Scientific), treated with DNase (#AM1907, Thermo Fisher Scientific) and reverse transcribed into cDNA (#4387406, Thermo Fisher Scientific). cDNA was diluted 1:1 in nuclease-free water and used to amplify *F. hepatica h2b* (*fhh2b*) with the FastStart Taq DNA Polymerase, dNTPack (#4738381001, Millipore Sigma) and 0.4 µM of each primer (S1 Table). A bacterial neomycin phosphotransferase gene (U55762) from the pEGFP-NI vector (Clontech) was used to generate control dsRNA. T7 promoter sequences were used on opposing primers to generate templates with T7 promoter sequences for subsequent RNA transcription. PCR product sizes were confirmed on a 1-2% agarose gel, before products were cleaned using the ChargeSwitch PCR Clean-Up Kit (#CS12000, Thermo Fisher Scientific). Sequences were confirmed via sequencing of cleaned PCR products (Eurofins Genomics). Clean T7 templates were used to generate dsRNA with the T7 RiboMAX Express RNAi System (#P1700, Promega). Resulting dsRNA was resuspended in nuclease-free water and quantified on a Denovix DS-11.

Following excystment, NEJs/juveniles were soaked in 100 ng/µL dsRNA (diluted in RPMI) for 24 hours twice a week for the duration of the experiment to ensure knockdown [84]. Worms were imaged each week to monitor growth using ImageJ (as described in [91]), and a subset of worms exposed to EdU to monitor proliferation, as described previously. At the conclusion of trials worms were snap frozen in liquid nitrogen with cDNA extracted as described above. Quantitative PCRs (qPCR) were run to assess target transcript knockdown in 10 µL volumes with 0.2 µM of each primer (S1 Table), 2 µL 1:1 diluted cDNA and SensiFAST SYBR No-ROX Kit (#BIO-98005, Bioline). Knockdown was assessed using ΔΔCt analysis as described by Pfaffl [92].

### Graphs and Statistics

Graphs and statistical tests were carried out in Graphpad Prism (v8, La Jolla, CA, USA). Data were tested for normality. Where data were normally distributed, parametric tests (t test or ANOVA) were used, with non-parametric tests (Mann-Whitney or Kruskal Wallis) used when data were non-normally distributed. Post-hoc analyses (Dunn’s, Tukey’s or Dunnett’s multiple comparisons tests) were employed to identify differences between multiple groups. Volcano plots were produced using R and heatmaps were produced in R (see GitHub code), or with the Broad Institute’s Morpheus (https://software.broadinstitute.org/morpheus). Unless otherwise stated mean ±SEM is used for graphs and in figures. Schematic figures were created with <u>BioRender.com</u>.

## Acknowledgements

Thanks to Dr Niall Byrne at Queen’s University Belfast for irradiating transcriptomic samples.

## Supporting Information

**S1 Fig. *F. hepatica* neoblast-like cells display temporal morphology changes.** (A) Images of juvenile *F. hepatica* cultured for three or ten days in CS50 growth inducing media with labelling of proliferative cells (EdU, green) and all nuclei (DAPI, purple); scale bars = 50 µm. (B) Comparison of EdU^+^ nuclei and EdU^-^ nuclei area in different aged juveniles shows that EdU^+^ cells are reduced in size by around half as worms age, whereas EdU^-^ cells increase slightly in size (Mann-Whitney U test). ****, p<0.0001.

**S2 Fig. Irradiation of *F. hepatica* juveniles significantly ablates their neoblast-like cells.** (A) Confocal images of juvenile *F. hepatica* that were cultured for three days *in vitro* before being dosed with 0 – 200 Gy radiation, then cultured for a further 72 h (final 24 h in EdU) and stained for EdU (green) shows that radiation ablates EdU^+^ nuclei; DAPI (magenta), scale bars = 50 µm. (B) # EdU^+^ nuclei in juvenile *F. hepatica* that were cultured for three days *in vitro* before being dosed with 0 – 200 Gy radiation shows significant reduction in number of EdU^+^ nuclei at 100 – 200 Gy (Mann-Whitney U test). ****, p<0.0001.

**S3 Fig. Only a subset of *F. hepatica* neoblast-like cells labelled with EdU prior to irradiation are susceptible to ablation.** (A) Timeline of experiment showing EdU incubation/labelling prior to irradiation. (B) Confocal images of *in vitro* juvenile *F. hepatica* treated according to timeline outlined above shows that cells already labelled with EdU (green) are not completely ablated by 200 Gy of radiation; DAPI counterstain (magenta). (C) # EdU^+^ nuclei in juvenile *F. hepatica* treated according to timeline outlined above shows that a significant number of EdU nuclei labelled prior to irradiation are ablated, though not all (Unpaired t test). **, p<0.01.

**S4 Fig. Genes downregulated in juvenile *F. hepatica* by irradiation are largely associated with neoblasts in *S. mansoni*.** (A) – Volcano plot showing differentially expressed genes (red) following irradiation of juvenile *F. hepatica* where blue shading indicates downregulated genes likely associated with the ablated neoblast-like genes. (B) Heatmap of expression (max z score, red; min z score, blue) in control and irradiated juvenile *F. hepatica* for the genes that are homologues of downregulated genes in adult *S. mansoni* following irradiation. (C) Venn diagram showing occurrence of homologues of *F. hepatica* downregulated genes following irradiation in various *S. mansoni* bioinformatic resources: downregulated in irradiated *S. mansoni* adults; enriched in adult stem cells (single cell); enriched in schistosomule stem cells (single cell).

**S1 Table. Primers used in PCR reactions.**

**S2 Table. DESeq2 results following irradiation of juvenile *Fasciola hepatica*.** Sheet 1 – Downregulated genes (adjp < 0.05), Sheet 2 – Upregulated genes (adjp < 0.05), Sheet 3 – All DESeq2 results.

**S3 Table. Transcripts downregulated in both *Fasciola hepatica* juveniles and *Schistosoma mansoni* adults following irradiation.**

**S4 Table. Transcript per million reads of all *Fasciola hepatica* transcripts identified during RNAseq.**

**S1 File. Nucleotide sequences of ‘novel’ MSTRG *Fasciola hepatica* transcripts identified by Stringtie during transcriptomic processing.**

